# Coral microbiomes from the Atlantic and Indo-Pacific oceans have the same alpha diversity but different composition

**DOI:** 10.1101/2024.08.16.608269

**Authors:** Maria E. A. Santos, James D. Reimer, Bogdan Kiriukhin, Hin Boo Wee, Masaru Mizuyama, Hiroki Kise, Marcelo V. Kitahara, Akira Iguchi, ‘Ale’alani Dudoit, Robert J. Toonen, Filip Husnik

**Author notes:** & (Maria E. A. Santos, Filip Husnik). Coral Biogeochemistry Laboratory, University of Hong Kong, Hong Kong (Maria E. A. Santos).

## Abstract

Corals are early-branching animals highly reliant on diverse symbionts for growth and reproduction. Most coral groups, including stony corals and hydrocorals, exhibit deep genetic divergence between the Atlantic (ATO) and Indo-Pacific (IPO) oceans, hampering their direct comparison. Although sibling zoanthid species (Hexacorallia: Zoantharia) deviate from this pattern, their symbioses have so far only been studied on local scales. Here, we examined the microbiomes of *Palythoa caribaeorum* from the ATO and *P. tuberculosa* from the IPO. Our extensive geographical sampling and metabarcoding revealed that *Palythoa* microbiomes have similar alpha diversity in both oceans. The primary exceptions are the symbiodiniacean *Cladocopium* and Chlamydiae bacteria, which mirror the global diversity patterns of corals. Despite distinct overall microbial compositions between oceans, some regions shared remarkably similar communities, hinting at the importance of both symbiont phylogeny and function. Finally, we explore the shift from commensal/mutualistic microbes to opportunistic pathogens, crucial amid the ongoing environmental changes.

## Introduction

Coral reefs are among the most diverse and valuable ecosystems that provide numerous resources such as fisheries, tourism, and bioprospection^1^. Most corals strongly depend on microbial symbionts for their growth and reproduction^2^. Due to this microbiome reliance, ecological importance, and phylogenetic position among early-branching animals, corals are an excellent and widely studied marine symbiosis model. Among the main threats to coral reefs is climate change as increasing seawater temperatures affect the fitness of both the coral animal and its associated microorganisms^3,4^. However, the global diversity patterns of coral microbiomes and how they respond to environmental stress are still far from being comprehensively understood^5^.

Corals feed by capturing prey organisms and absorbing organic matter from the water column^6,7^, but a part of their nutritional requirements in shallow waters is met by biosynthetic products of their photosynthetic symbionts. The photosymbiotic eukaryotes are dinoflagellates from the family Symbiodiniaceae, known as symbiodiniaceans. Although coral-dinoflagellate symbiosis was detected since the Late Triassic^8^, the relationship with symbiodiniaceans was predicted to have originated in the early to mid-Jurassic period^9^. Nevertheless, in most coral species, the symbionts are horizontally acquired every generation from the environment^10^. Each coral colony usually hosts a community of Symbiodiniaceae lineages that can change over time depending on the host and environmental conditions^11^. Consequently, distinct physiological capabilities were described for different symbiodiniacean clades and species^9^, and some corals can rapidly shuffle their symbiont community when stressed^12,13^. However, extreme changes in seawater conditions, such as temperature or pH, can lead to the expulsion of the symbiodiniacean cells, a phenomenon known as coral bleaching, and eventually result in coral death^3,14^.

Aside from Symbiodiniaceae, very little is known about the role of other protists on coral biology^5,15^. Among understudied coral-associated eukaryotes, corallicolids^16,17^, chromerids^18^, ciliates^19^, and several lineages of fungi^20,21^ are frequently reported worldwide. Corals also house diverse prokaryotic symbionts. The mutualistic bacterial community of corals supports the nutrient cycling of their host, such as nitrogen^22^ and sulfur^23^, and provides resistance against microbial pathogens^24^. Unfortunately, our understanding of the distribution of coral-associated bacteria across large spatial scales is still limited, which hampers ongoing efforts to identify probiotic candidates to improve coral resilience in locations likely to be most affected by the rising temperature^25,26^. Even less is known about coral-associated Archaea, which were proposed to play a role in the holobiont nitrogen recycling^27^.

Following global diversity patterns in most terrestrial and aquatic ecosystems^28^, the diversity in coral reefs increases from higher latitudes towards the equator. Reefs in the Indo-Pacific Ocean (IPO) have a higher diversity than in the Atlantic Ocean (ATO)^29,30,31^, partly due to IPO’s larger area, older geological age, and lower extinction rates. The Central Indo-Pacific, also known as the Coral Triangle, is a major marine biodiversity hotspot. Although several non-exclusive hypotheses have been proposed to explain this species’ richness pattern^32,33^, these studies have focused on the diversity of animals and macroalgae. Understanding if and how the coral host shapes the diversity of associated microbes will contribute to forecasting the future of reefs in the face of ongoing human disruptions to marine ecosystems.

Most coral families have a deep genetic divergence among species of the IPO and ATO, including stony corals^34^ and hydrocorals^35^. Zoanthids (Anthozoa: Hexacorallia: Zoantharia) are an exception to this pattern, with sister species between the IPO and ATO^36,37^. Strikingly, some of the sibling pairs might even be species complexes with global distribution^38^. In this study, we focused on the most widespread zoanthid sibling species *Palythoa caribaeorum* which has an amphi-Atlantic distribution^39^, and its sibling *P. tuberculosa*, which is widely distributed in the IPO^40^.

Zoanthids are emerging as a unique model system to examine broad patterns of symbiosis in the same coral lineage. Since the *Palythoa* microbiome is most likely acquired horizontally from the environment^41^, fundamental questions such as what acts as the unit of selection in holobionts can be addressed in this system to distinguish whether it is the biological functions (such as specific metabolites) as proposed by the “It’s the song, <u>not</u> the singer (ITS<u>N</u>TS)” hypothesis^42^ or both biological functions and taxa as reported in the “It’s the song <u>and</u> the singers (ITS<u>A</u>TS)” response to the hypothesis^43^. Another rarely addressed fascinating question is how hostassociated microbial communities assemble in marine environments under different geographic and environmental conditions^44,45^. Surprisingly, zoanthid symbioses have been so far investigated only at small to regional scales^46–48^. Here, we bridged this gap and revealed global patterns in the diversity of prokaryotes and microeukaryotes associated with *P. caribaeorum* and *P. tuberculosa*. Using the most extensive geographical sampling for such a marine symbiosis system to date, we examine the entire microbiome of *Palythoa* and discuss the diversity and evolution of symbiotic interactions across the ATO and IPO (Figure 1).

**Figure 1:**
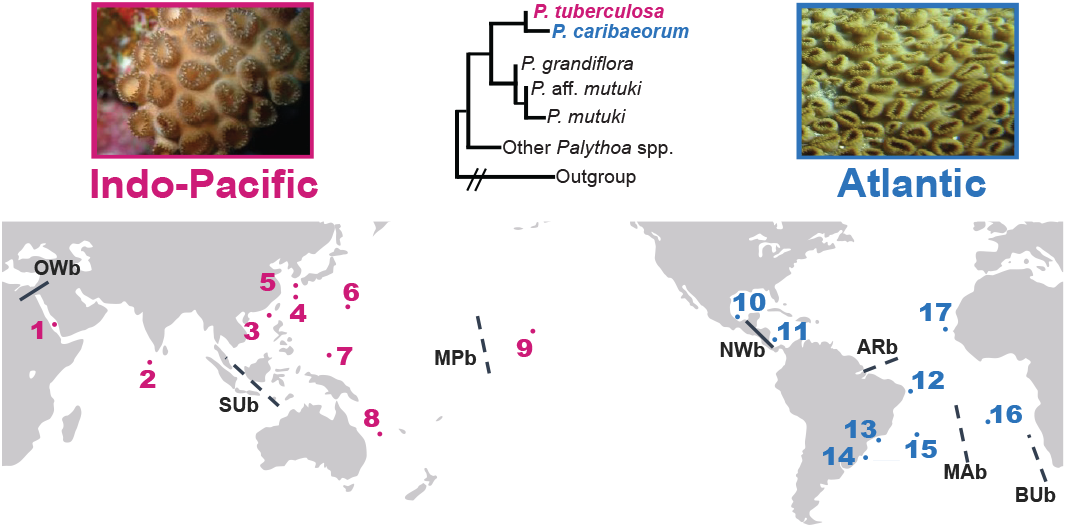
Sampling regions and main global biogeographical barriers. Sampling campaigns in the Indo-Pacific Ocean (*P. tuberculosa*, in pink) were conducted in the 1-Red Sea, 2-Maldives, Taiwan (3-Dongsha Atoll), Japan (4-Okinawa Island, 5-Iotorishima Island, and 6-Ogasawara Islands), 7-Palau, 8-New Caledonia, and 9-Hawai’i, while in the Atlantic Ocean (*Palythoa caribaeorum*, in blue), samples were collected in the Caribbean Sea (10-Mexico and 11-Costa Rica), Brazil (12-Rocas Atoll, 13-Rio de Janeiro, 14-Xavier Island, and 15-Trindade Island), 16-Saint Helena Island, and 17-São Vicente Island. On a global scale, the hard biogeographical barriers between the Indo-Pacific and Atlantic oceans are the final closure of the Isthmus of Panama (New World barrier; NWb) and the Tethys Seaway (Old World barrier; OWb). On ocean basin scales, there are five main soft barriers: Amazon-Orinoco rivers (ARb), Benguela upwelling system (Bub); Sunda Shelf (Sub), and the large open ocean distances Mid-Pacific (MPb) and Mid-Atlantic (MAb). The schematic phylogenetic tree is based on the literature^36^.

## Results

### Global diversity of prokaryotes and eukaryotes associated with *Palythoa* corals

Our results revealed 25,816 ASVs of prokaryotes and 5,781 ASVs of eukaryotes (ESM Tables 2, and 3). Based on this taxonomic assignment, *Palythoa* coral samples were associated with 30 bacterial phyla, one archaeal phylum, and 13 main lineages of eukaryotes (Figure 2). Almost all prokaryotes were identified as bacteria. While two *Palythoa* samples were associated with Thaumarchaeota, they contained less than 5 read pairs each (Figure 2A; ESM Table 2). Globally, the most common bacteria associated with *Palythoa* were from the phylum Proteobacteria (synonym Pseudomonadota; Figure 2A, ESM Table 5). At the class level, Gammaproteobacteria was the most common, followed by Alphaproteobacteria, Flavobacteriia, and Cyanobacteria. Most bacterial genera were similar between the ATO and the IPO, with *Endozoicomonas* and *Vibrio* being the most common worldwide (ESM Table 5).

**Figure 2:**
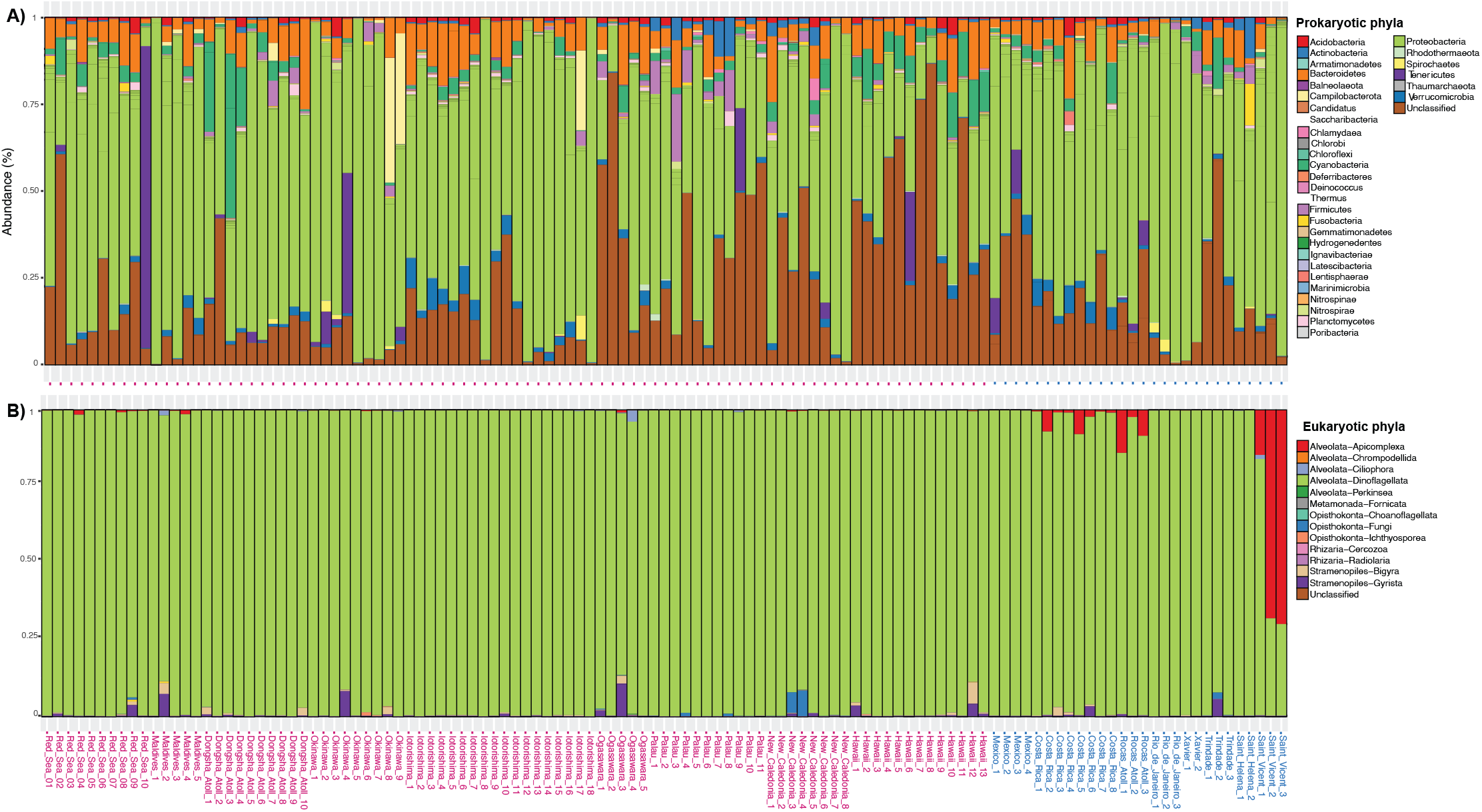
Relative abundance (%) of prokaryotic (A) and eukaryotic (B) phyla associated with each *Palythoa* sample. The abundance of prokaryotes and microeukaryotes was estimated using MiSeq sequencing of 16S rRNA and 18S rRNA gene amplicons. Coral samples from the Indo-Pacific Ocean (*P. tuberculosa*, in pink), samples were collected in the Red Sea, Maldives, Taiwan (Dongsha Atoll), Japan (Okinawa Island, Iotorishima Island, and Ogasawara Islands), Palau, New Caledonia, and Hawai’i, while in the Atlantic Ocean (*P. caribaeorum*, in blue) were collected in the Caribbean Sea (Mexico and Costa Rica), Brazil (Rocas Atoll, Rio de Janeiro, Xavier Island, and Trindade Island), Saint Helena Island, and São Vicente Island.

As expected, dinoflagellates dominated the eukaryotic reads of all colonies, except two *Palythoa* samples that had a higher percentage of apicomplexan reads (Figure 2B). Apicomplexa was the second most common lineage of eukaryotes associated with *Palythoa* (ESM Table 5), while other commonly observed groups were Rhodophyta, Gyrista (mainly diatoms), Fungi, Ciliophora, and Chlorophyta. At the family level, Symbiodiniaceae were dominant in most samples, and an additional four families were observed as the most common globally: apicomplexan of Corallicolidae, rhodophytes of Corallinales and Gigartinales, and fungi of Exobasidiomycetes. While the prokaryotic data showed that the most common genera are largely shared between the ATO and IPO, an opposite pattern was observed from the eukaryotic data where the most common genera observed at each ocean basin were distinct (ESM Table 5).

Within the family Symbiodiniaceae, reads were classified into 518 ‘Defining Intragenomic Variants’ (DIVs) and four genera (Figure 3A and ESM Table 4). *Cladocopium* was associated with all samples, followed by DIVs of *Durisdinium, Symbiodinium*, and *Brevolium* (total of 402, 65, 48, and 3 DIVs). Two *Cladocopium* ‘type-profiles’ were observed globally (C1-C1c-C1b and C3-C1-C42.2), while six and sixteen were reported only to AO and IPO, respectively (Figure 3B). The DIVs with the highest number of reads across both oceans were C1, C3, and C1n (Figure 3C).

**Figure 3:**
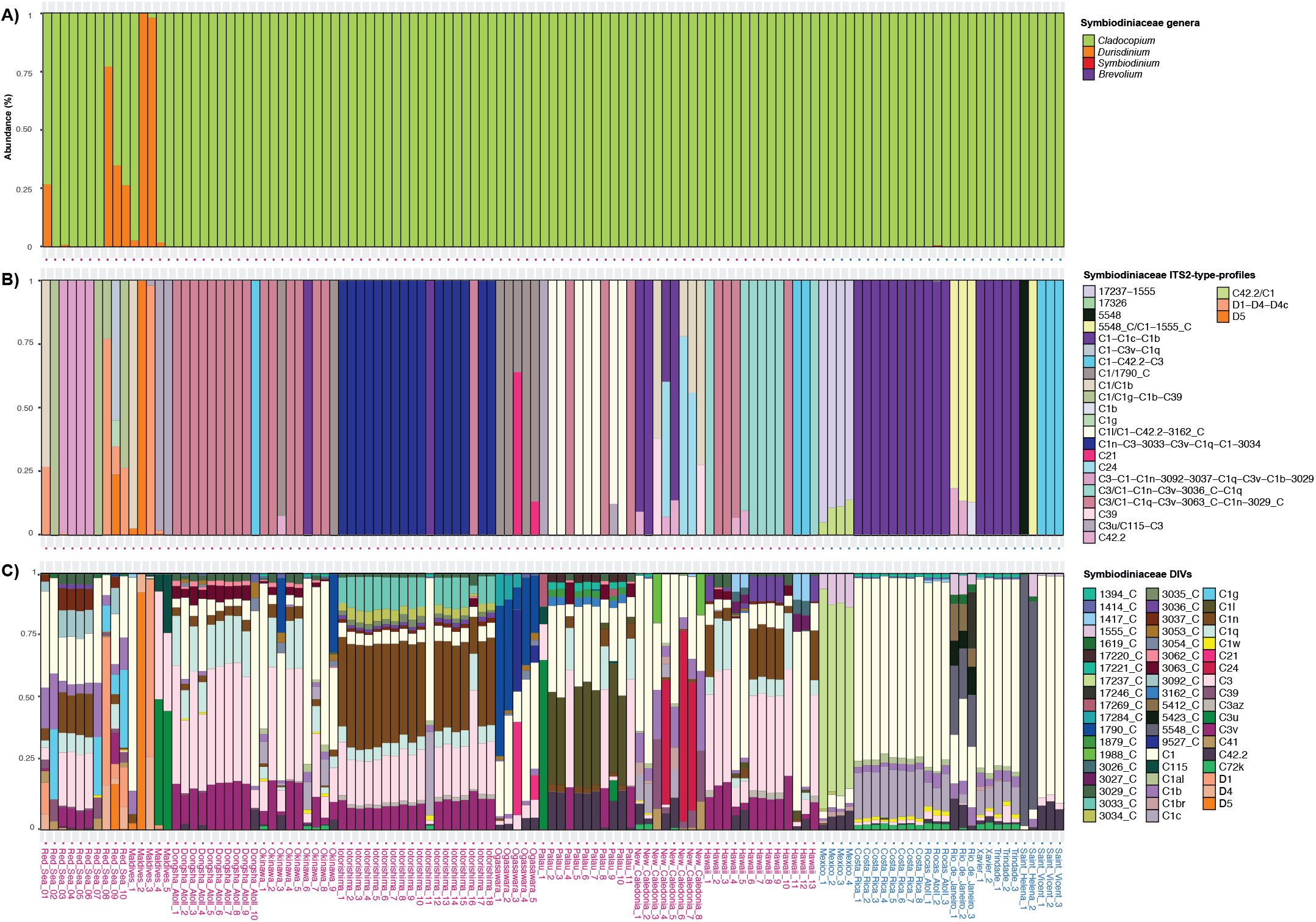
Relative abundance (%) of Symbiodiniaceae associated with each *Palythoa* sample by genera (A), ‘ITS2-type-profiles’ (B), and ‘Defining Intragenomic Variants’ (DIVs; C). Symbiodiniaceae abundance was estimated using MiSeq sequencing of the ITS2 rRNA spacer amplicon. Coral samples from the Indo-Pacific Ocean (*P. tuberculosa*, in pink), samples were collected in the Red Sea, Maldives, Taiwan (Dongsha Atoll), Japan (Okinawa Island, Iotorishima Island, and Ogasawara Islands), Palau, New Caledonia, and Hawai’i, while in the Atlantic Ocean (*P. caribaeorum*, in blue) were collected in the Caribbean Sea (Mexico and Costa Rica), Brazil (Rocas Atoll, Rio de Janeiro, Xavier Island, and Trindade Island), Saint Helena Island, and São Vicente Island. Only the most abundant DIVs are displayed, and labels indicate the Symportal^64^ database ID number followed by the genus the sequence is from (C=*Cladocopium* and D=*Durusdinium*).

Within prokaryotes, core microbes included the gammaproteobacterial genera *Vibrio* (99%), *Endozoicomonas* (87%), *Tenacibaculum* (83%), *Photobacterium* (82%), and *Alteronomas* (79%), and the alphaproteobacterial genus *Ruegeria* (78%) (Figure 2A; ESM Table 2). Among eukaryotes, only the symbiodiniacean *Cladocopium* was detected in over 75% of *Palythoa* samples (Figures 2 and 3; ESM Tables 3 and 4).

### *Palythoa* microbiomes from the Atlantic and Indo-Pacific oceans have the same alpha diversity but different composition

The prokaryotes and eukaryotes associated with *Palythoa* had a similar Shannon index diversity in both the ATO and IPO (*p* = 0.55 and 0.28, respectively; Figure 4A-B). Within prokaryotes, the only statistical difference was observed for the phylum Chlamydiae, which was more abundant in samples from the IPO (*p*=0.03; Figure 5A; ESM Table 8). Within eukaryotes, the overall diversity of symbiodiniaceans was higher in the IPO (Figure 5C; *p*=3×10^−11^) for both genera with the highest number of reads, *Cladocopium* (*p*=4×10^−9^) and *Durisdinium* (*p* = 0.02). Chromerids were mostly detected from the IPO, although with only 250 reads assigned and no statistical significance (Figure 5B; *p* = 0.12). On the other hand, corallicolids and fungi were significantly more associated with *Palythoa* in the ATO (*p*= 1.5×10^−6^ and *p* = 0.01, respectively).

**Figure 4:**
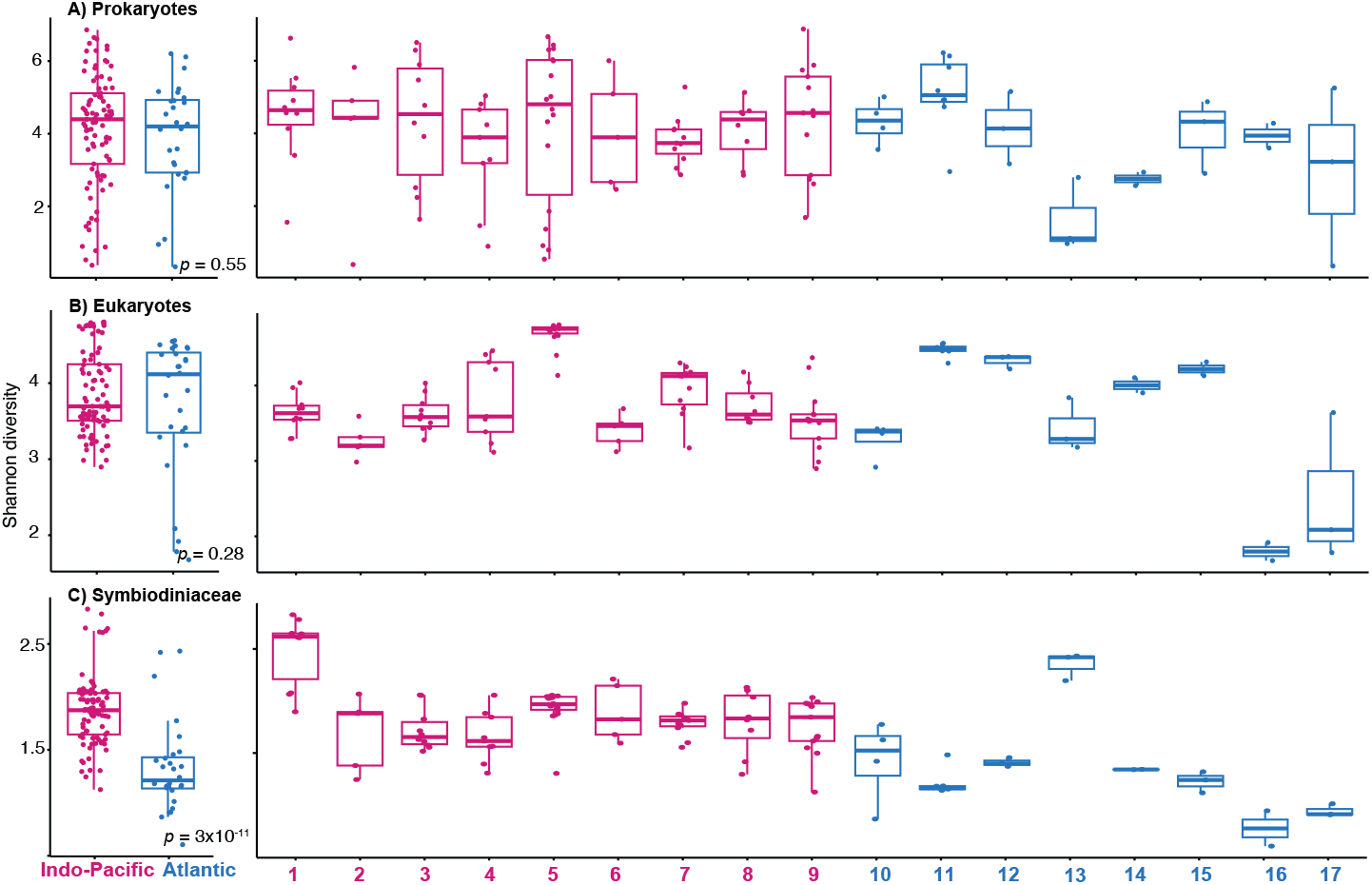
Shannon diversity index of *Palythoa*-associated microorganisms across each ocean (left) and region (right). The diversity values of prokaryotes (A), eukaryotes (B), and Symbiodiniaceae (C) were estimated using MiSeq sequencing of 16S rRNA, 18S rRNA, and ITS2 rRNA amplicons. Coral samples from the Indo-Pacific Ocean (*P. tuberculosa*, in pink) were collected in the 1-Red Sea, 2-Maldives, Taiwan (3-Dongsha Atoll), Japan (4-Okinawa Island, 5-Iotorishima Island, and 6-Ogasawara Islands), 7-Palau, 8-New Caledonia, and 9-Hawai’i, while in the the Atlantic Ocean (*P. caribaeorum*, in blue) samples were collected in the Caribbean Sea (10-Mexico and 11-Costa Rica), Brazil (12-Rocas Atoll, 13-Rio de Janeiro, 14-Xavier Island, and 15-Trindade Island), 16-Saint Helena Island, and 17-São Vicente Island.

**Figure 5:**
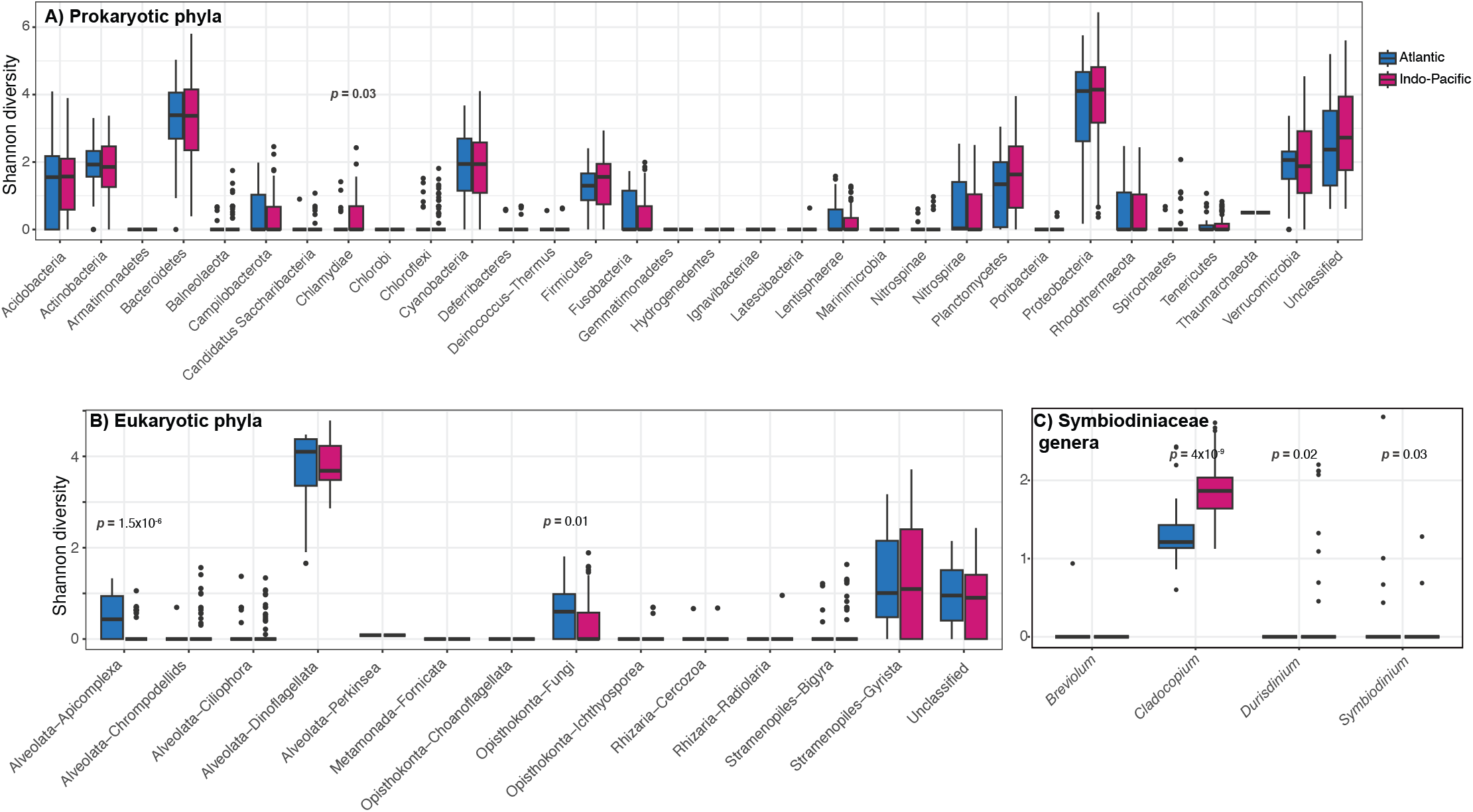
Shannon diversity index of *Palythoa*-associated microorganisms shown for phyla of prokaryotes and eukaryotes, and genera of Symbiodiniaceae. Taxa from the Atlantic and Indo-Pacific oceans are shown in left (blue) and right (pink) bars. The diversity values of prokaryotes (A), eukaryotes (B), and Symbiodiniaceae (C) were estimated using MiSeq sequencing of 16S rRNA, 18S rRNA, and ITS2 rRNA amplicons.

Despite a similar alpha diversity for most bacteria and eukaryotes, and the nMDS showing a mix of samples from ATO and IPO (Figure 6), the PERMANOVA and most regional post-hoc tests revealed an overall distinct symbiont composition for *Palythoa* from different ocean basins, regions within each ocean, and at different sea surface temperatures (ESM Table 6). Nevertheless, 19 pairwise tests showed a similar microbial composition between regions of different oceans (five pairs of regions for bacteria, another five for eukaryotes, and nine for Symbiodiniaceae; ESM Table 7). Regions from the AO had a higher number of pairwise tests showing a similar microbial composition in contrast to regions from the IPO (17, 17, and 15 compared to 0, 1, and 3).

**Figure 6:**
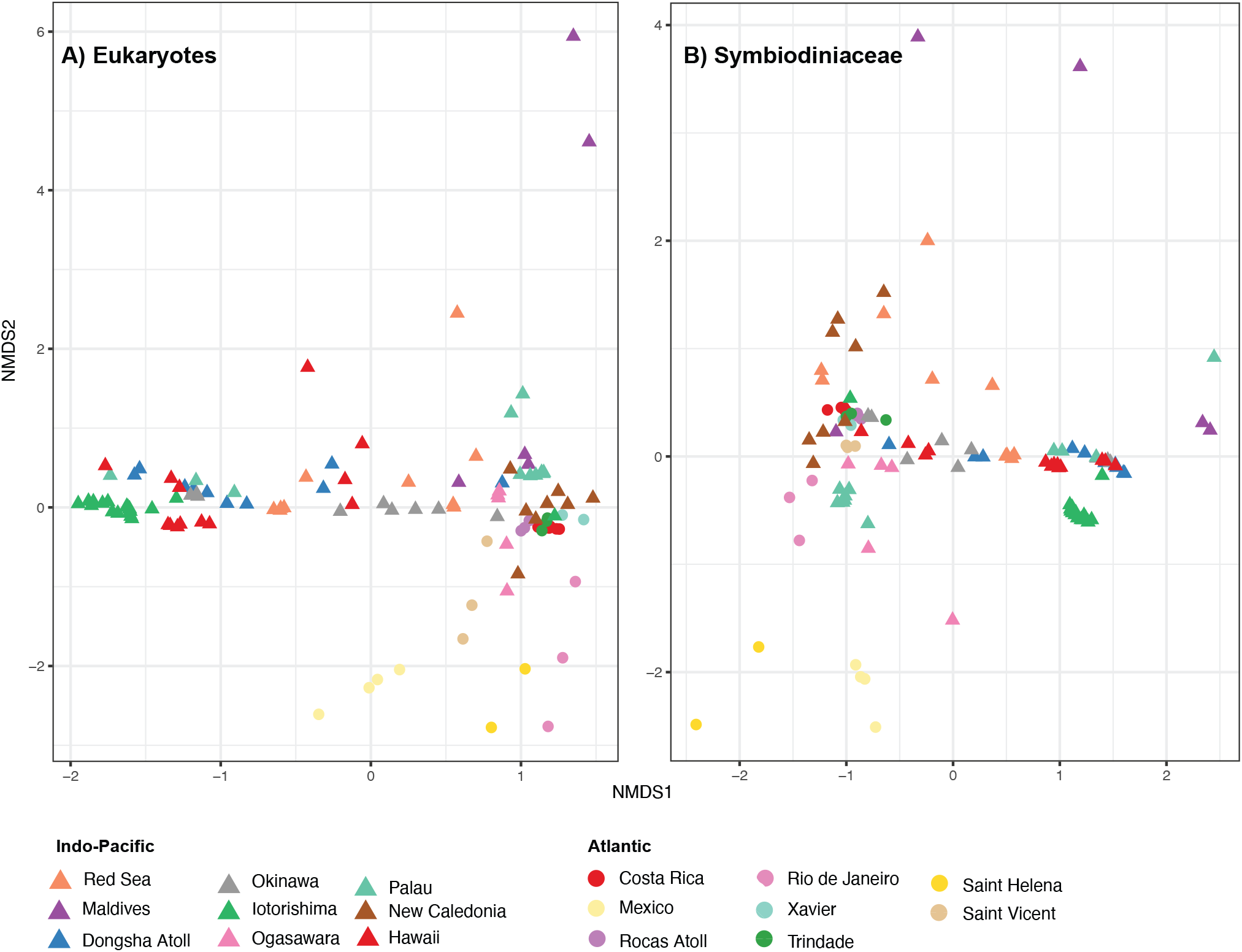
Non-metric multidimensional scaling analysis of the Bray-Curtis distances assessing the similarity of microbial composition across regions. Plots had a stress of 0.28 (not displayed; prokaryotes), 0.15 (A; eukaryotes), and 0.15 (B; Symbiodiniaceae). The diversity values were estimated using MiSeq sequencing of 16S rRNA, 18S rRNA, and ITS2 rRNA amplicons. Coral samples from the Atlantic Ocean (*Palythoa caribaeorum*) were collected in the Caribbean Sea (Mexico and Costa Rica), Brazil (Rocas Atoll, Rio de Janeiro, Xavier Island, and Trindade Island), Saint Helena Island, and São Vicente Island, while in the Indo-Pacific Ocean (*P. tuberculosa*) samples were collected in the Red Sea, Maldives, Taiwan (Dongsha Atoll), Japan (Okinawa Island, Iotorishima Island and Ogasawara Islands), Palau, New Caledonia, and Hawai’i.

## Discussion

### Sediment embedding in *Palythoa* and the potential role of associated organisms as epibionts, prey, and in toxin production

It has been proposed that sediment might act as a seed bank for bacteria to colonize stony coral surfaces, as coral mucus shares more bacteria with sediment than seawater^49^. Although most zoanthids, including *Palythoa* species, do not have a calcium carbonate skeleton, they incorporate sediment inside their tissues. Particles such as sand, sponge spicules, and multicellular algae can represent up to 45% of their wet tissue weight^50^. This strategy likely improves the internal structure of the colonies^50^ and makes the tissue less palatable to predators^51^. As microbes or their DNA are also likely to be incorporated inside the *Palythoa* tissue during this process, distinguishing the sediment-acquired component from truly host-associated organisms is challenging and we interpret any rare sequences with caution. For example, the diatom *Psammodictyon* and the dinoflagellate *Ostreopsis* were ranked among the most common eukaryotes recorded for *Palythoa* colonies (ESM Table 5). Still, these benthic microbes were not previously reported as coral symbionts and it is possible that such associations also occurred due to sediment incorporation (or prey) rather than symbiosis. Additionally, some bacteria might not be directly associated with *Palythoa* but instead symbiotic with symbiodiniaceans or other coral protists^52,53^.

*Ostreopsis* and *Palythoa* are the main organisms reported to contain one of the most potent marine neurotoxins, palytoxin^54–56^, however, there was no clear relationship reported between the two taxa. Palytoxin can be sequestered in the food web by fishes and crustaceans^57^ causing severe seafood poisoning^58,59^, as well as accidental contact with skin by hobbyists with contaminated aquaria waters^60,61^. Here, we detected six ASVs assigned to *Ostreopsis* in eight *Palythoa* samples from three locations in the ATO and ten samples from four locations in the IPO (926 reads and 54 reads; ESM Table 3). As *Ostreopsis* co-occurred with several bacterial lineages (ESM Table 9), our data point toward a complex relationship where diverse bacteria can be associated with toxin presence as reported previously^62^. Nonetheless, if *Ostreopsis* or their bacteria synthesize palytoxin, their presence in *Palythoa* tissues could explain palytoxin accumulation.

Our results showed taxa previously reported to bore skeletons of stony and hydrocorals^21,63^ including ASVs assigned as rhodophytes, *Phaeophila* green algae, Ascomycota and Basidiomycota fungi, and Gyrista (ESM Table 3), revealing that their niche is not restricted to endolithic environments. Other detected microeukaryotes are potentially opportunistic pathogens or accessory symbionts with an unclear effect on the host, for example, the ciliate *Ephelota* that was associated with two samples. The species *E. mammillata* was hypothesized to be a commensal epibiont that uses the host as a substrate to optimize feeding^64^, and we provide additional evidence for this occasional *Ephelota*-corals interaction.

### Non-photosynthetic microbial eukaryotes associated with

#### Palythoa

Not only symbiodiniaceans, but other diverse other microeukaryotes can influence coral health, still, our understanding of their global distribution is limited. Apicomplexa are well-known animal parasites but the relationship between Corallicolida and their host is unclear^65,66^. Corallicolids have been reported worldwide and are the second most common endosymbiotic protist associated with corals^16,66^. Remarkably, corallicolids (and fungi with lower support) were detected as more common in *Palythoa* samples from the ATO. Aquarium experiments simulating marine heat waves suggested that corallicolids increase their abundance in stressed octocoral colonies^64^, implying that their higher presence in *Palythoa* from the ATO might be stress-induced. Although we do not have data to confirm that these colonies were completely healthy, they were not positively associated with well-known pathogens such as *Vibrio* spp. or ciliates (ESM Table 9). While the coral tissue did not have any visible damage, corals from ATO could be more stressed without a visible difference. Clarifying the corallicolid dispersal mode and interactions with hosts across shallow and deep-sea waters will offer potential explanations for their association with Atlantic corals. Corallicolids likely have a complex life cycle similar to other apicomplexans^65,66^. For instance, a potential life stage could be associated with reef fishes, as fish’ feces are an important transmission mode of symbiodiniaceans^67^, and the sister group of corallicolids also infects fish^68^.

Among fungi, we found eight families associated with *Palythoa* (ESM Table 3). Many of these lineages are better known from terrestrial environments raising a question whether these fungi represent contamination from wind-born spores or freshwater run-offs. However, crossing the salt barrier is known as an important aspect of fungal diversification^69^. Sequences of ‘terrestrial’ fungi were previously found in stony corals and hydrocorals^20,70^ and reef-associated algae^71^, implying that at least some of the fungi detected here from *Palythoa* are native members of its microbiome. Coral-fungal interactions are still poorly understood, and observations range from opportunistic pathogens^72^ to potential mutualists^73^.

The relationship between ciliates and corals is also multifaceted. Ciliates play an important role as bacterial predators in the coral mucus, removing populations of pathogenic bacteria and assisting with nutrient cycling^74^. In contrast, several ciliated species were reported associated with the black, brown, and white band diseases^19^. *Holosticha diademata*, one of the most common ciliate causatives of diseases, was not found associated with *Palythoa* in the assigned ASVs, but our data included five other genera of coral ciliates, as well as *Chlamydonellopsis* that was not previously reported with corals.

### Walking a fine line between mutualism and pathogenicity in coral-symbiont interactions

Certain coral-associated microorganisms are known to be capable of acting as mutualists under some conditions and as pathogens under others^75^. How the overall microbiome diversity and environmental factors contribute to these transitions is, however, often limited by our understanding of a healthy microbiome baseline for a particular location. The bacterial genus *Endozoicomonas* is widely presumed to be the main mutualist of corals and other cnidarians^76^. Nevertheless, this perception of pure mutualism was recently challenged, suggesting that a more cautious and case-specific interpretation is needed and that the position of diverse *Endozoicomonas* lineages on the symbiosis spectrum is likely more varied and context-dependent^77^. On the other hand, Vibrionaceae, and especially the genus *Vibrio*, are usually portrayed only as opportunistic pathogens. Vibrionaceae were reported to cause tissue loss in diseased corals worldwide^78,79^ and bleaching of *Acropora* stony corals was shown to lead to a shift from colonies dominated by *Endozoicomonas*-allied to *Vibrio*-allied bacteria^80^. What is the role of Vibrionaceae in the healthy microbiome and the exact trigger of this pathogenicity nonetheless remains unclear.

In our data, *Palythoa* colonies associated with *Vibrio* usually had low numbers of *Endozoicomonas*, although no statistical difference was observed (ESM Table 9). Unexpectedly, the co-occurrence of *Endozoicomonas* and the corallicolid *Anthozoaphila* was significantly positively correlated. Still, additional research is needed to examine their potential interaction. Notably, Tenericutes was the fourth most common phylum in the IPO and the fifth in the ATO (Figure 2A and ESM Table 2). These ASVs mostly come from the genera *Spiroplasma* and *Mycoplasma* which are known as animal pathogens. However, several lineages of these marine Tenericutes were also proposed to be potential defensive mutualists of invertebrates, including corals^81^. Our data support the notion that relationships among members of the coral holobiont are intertwined, and further experiments are needed to clarify the environmental and microbiome diversity tipping point that shifts some microbiome members toward pathogenicity.

### Strongly host-associated microbes mirror the diversity pattern of corals

The most surprising result of our study is that, unlike the classic pattern in marine biogeography that the IPO has a much higher diversity than the ATO, microbes associated with corals have a similar alpha diversity between the oceans (Figure 4). Only symbiodiniaceans, corallicolids, fungi, and chlamydial bacteria were exceptions and had a significantly different diversity between the oceans. Symbiodiniaceae showed higher diversity in the IPO and followed the biogeographic patterns of other reef organisms^29,32^ (Figure 5). No other coral-associated eukaryotes were detected as more diverse in the IPO. However, the generally low number of non-Symbiodiniaceae reads could lead to false negatives in lineages such as chrompodelids (ESM Tables 3 and 8). Conversely, our results revealed the opposite trend for corallicolids and fungi that were significantly more diverse in *Palythoa* samples from the ATO.

Among prokaryotic phyla, Chlamydiae showed higher diversity in the IPO. Chlamydiae are well-known endosymbionts of diverse animals and unicellular eukaryotes, and the assigned ASVs belong to *Simkania* (ESM Table 2), previously reported as coral-associated and forming aggregates with *Endozoicomonas*^82^. Symbiodiniaceae were recently also shown to house intracellular bacterial communities^52^ including Chlamydiae^83^. Unless free-living and sediment associated, Chlamydiae detected in *Palythoa* are most likely either coral symbionts, dinoflagellate symbionts, or capable of infecting both multicellular and unicellular hosts. This result implies a nested pattern of symbiotic diversity where the higher diversity of corals in the IPO potentially drives not only a higher diversity of their symbiodiniaceans but also of bacterial endosymbionts of both dinoflagellates and corals.

### On a global scale, symbiodiniaceans and a few bacterial lineages are core members of the coral microbiome

That symbiodiniaceans are integral symbionts of most corals is rarely questioned. However, which other microorganisms (if any) are the core members of coral microbiomes globally is a long-standing coral microbiome question^84–86^. Previous studies have mostly investigated global diversity patterns of Symbiodiniaceae using samples from diverse stony coral species^87,88^. We opted for an alternative approach and analyzed a pair of closely related species to reduce the effects of the host specificity and examine fine-scale microbiome diversity worldwide. *Palythoa* likely rely on the environmental acquisition of symbionts^41^, which increases the frequency of symbiont replacements and/or co-symbioses. Given the high potential of *Palythoa* to rely on heterotrophy^84,85^ compared to other coral species that rely more on autotrophic photosynthetic products of symbiodiniaceans^4^, it is surprising that *Palythoa* had a high level of specificity for *Cladocopium* as the key symbiont selected in most populations (Figure 3). Symbiodiniaceae diversity at the level of type-profiles and DIVs revealed that the community was shaped both by region and seawater temperature (Figure 6, ESM Table 6). Corals that have a high heterotrophy feeding performance, including *P. tuberculosa*, have higher resilience during bleaching events, providing them with a competitive advantage under climate change-induced stress^4,7,89^. If and how the different *Cladocopium* profiles and DIVs contribute to the *Palythoa* generalists being able to outcompete other corals during stress remains to be experimentally tested. With ongoing climate changes, the relationship context is likely to change. Some symbiodiniaceans might switch from commensal or mutualistic into a more parasitic relationship, which may trigger other changes in the symbiont composition^75^. Our study lays a foundation for close monitoring of how the distribution of symbiodiniaceans will change in the upcoming decades on a large geographical scale.

The bacterial component of coral microbiomes is less stable than symbiodiniaceans and highly dependent on the environmental context, which often leads to contradictory results^81^. Our results from *Palythoa* (Figure 2) are congruent with previous reports on the prokaryotic diversity associated with other corals which showed that the bacterial genera *Endozoicomonas, Vibrio*, and *Rugeria* are associated with the greatest number of species^84,85^. Remarkably, several regions between oceans had a similar composition of *Palythoa-*associated organisms, and regions within AO shared more similar microbe communities, compared to IPO. Further research on the population genomics of zoanthids is crucial to elucidate how biogeographical barriers and filters influence the connectivity of host genotypes and their associated symbionts across the globe.

### Do marine animals select symbionts from seawater based on both the song and the singer?

Two central hypotheses have been proposed to explain the unit of symbiont selection by hosts: 1) biological functions (i.e., metabolites) as proposed by the “It’s the song, <u>not</u> the singer” hypothesis^42^; or 2) both biological functions and taxa as reported in the “It’s the song <u>and</u> the singers”^43^. The composition and abundance of the community associated with corals are distinct from the surrounding environment^49,84^. That *Palythoa* associate with microorganisms from the same clades across the globe suggests that in this marine symbiosis, the ‘singer’ matters and is selected surprisingly specifically given the predicted horizontal acquisition and low dependence on symbiont nutrients. What is the contribution of ‘songs/functions’^90^ for the selection specificity remains to be investigated. The future steps for *Palythoa* microbiome research should thus be generating genomes and transcriptomes of the main microbes to understand their phylogeny and functions (“songs”), and untangling interactions of diverse members of the coral holobiont. Finally, deciphering the transition of microbes from commensal/mutualistic to opportunistic pathogens will be essential for the future of coral reefs under climate change.

## Methods

### Sampling

Small fragments of colonies of *Palythoa tuberculosa* and *P. caribaeorum* were collected using snorkeling and SCUBA diving in depths ranging from 0 to 30 meters. Samples were collected across 17 regions between 2011 and 2022 (Figure 1, ESM Table 1). *Palythoa* species are not CITES protected. Tissue samples were fixed in 99% EtOH or frozen after collection. Sampling campaigns in the Indo-Pacific Ocean (*P. tuberculosa*) were conducted in the 1-Red Sea, 2-Maldives, Taiwan (3-Dongsha Atoll), Japan (4-Okinawa Island, 5-Iotorishima Island and 6-Ogasawara Islands), 7-Palau, 8-New Caledonia, and 9-Hawai’i. In the Atlantic Ocean (*P. caribaeorum*), samples were collected in the Caribbean Sea (10-Mexico and 11-Costa Rica), Brazil (12-Rocas Atoll, 13-Rio de Janeiro, 14-Xavier Island, and 15-Trindade Island), 16-Saint Helena Island, and 17-São Vicente Island. Although this sampling strategy allowed us to compare the microbiome of close-related hexacorals globally, it was not possible to sample in the same year/season worldwide due to time, and logistics. It should also be noted that the colonies of the coral samples are of different ages, and at different distances from each other due to site-specific differences across the *Palythoa* sampling sites. Whenever possible, we analyzed at least three samples per region, except for locations 14, 16, and 17 which only had two samples analyzed. In total, 117 samples were used for DNA extraction and amplicon sequencing. Tissue samples were deposited in the Fujukan Museum at the University of the Ryukyus, Okinawa, Japan (ESM Table 1). Approximate average surface seawater temperature was acquired from Giovani NOAA for the previous five years of the sampling date at each region (ESM Table 1).

### DNA extraction, library preparation, and Illumina MiSeq sequencing

After rinsing with distilled water and dissecting the *Palythoa* colonies, small fragments of the full polyps, including the gastric cavity, tentacles, and polyp wall were used for the DNA extraction. The total genomic DNA was extracted using one of the following kits: Qiagen DNEasy Blood & Tissue, Qiagen DNEasy PowerSoil, New England Biolabs Monarch Nucleic Acid Purification, or Lucigen MasterPure Complete DNA & RNA Purification, following the manufacturer’s instructions (ESM Table 1). The concentration of the DNA samples was measure using the Qubit Fluorometer with the Qubit 1X dsDNA HS Assay kit (Invitrogen, Japan). For amplicon sequencing of prokaryotes and microeukaryotes, we used PCR amplification with primers that target their 16S rRNA and 18S rRNA gene regions, respectively. As Symbiodiniaceae is by far the most abundant lineage of microeukaryotes associated with corals, a primer targeting their second internal transcribed spacer (ITS2 rRNA) region was also used to identify their diversity at higher resolution.

The V6-V8 region of the archaeal and bacterial 16S rRNA gene was amplified using the primers B969F 5’-ACGC GHNRAACCTTACC-3’ and BA1406R 5’-ACGGGCRGTGWGTRCAA-3’^91^. The V4 region of the 18S rRNA gene was amplified with a two-step amplification using the non-metazoan primer pairs EUK581-F 5’-GTGCCAGCAGCCGCG-3 with EUK1134-R 5’-TTTAAGTTTCAGCCTTGCG-3’, and E572F 5’-CYGCGGTAATTCCAGCTC-3’ with E1009R 5’-CRAAGAYGATYAGATACC RT-3’^92^. The ITS2 Rrna spacer was amplified with the primers Dino-forward 5’-TCGTCGGCAGCGTCAGATGTGTATAA-GAACAGGTGAATTGCAGAACTCCGTG-3’its2rev2-reverse GTCTCGTGGGTCGGAGATGTGTATAAGAGA- and 5’-CAGCCTCCGCTTACTTATATGCTT-3’^93^. The amplicon libraries were pooled and sequenced on four lanes of the MiSeq Illumina platform using paired-end reads (2x 300 bp). Detailed protocols are provided as ESM Protocol 1.

### Sequence data processing and statistical analyses

All raw reads are available under the NCBI BioProject accession number (XXXXX) and the full bioinformatics pipeline with all scripts is available from the Github repositories https://github.com/SantosMEA and https://github.com/ECBSU. For reads of the 16S rRNA and 18S rRNA gene amplicons, primers were removed using Cutadapt v4.1^94^, and further data processing and analyses were performed in R using the DADA2 pipeline^95^. Reads were denoised and joined into amplicon sequence variants (ASVs). Chimeras were removed using the command “removeBimeraDenovo” with the method “consensus”. The taxonomic assignment of 16S rRNA ASVs (prokaryotes) was done using RDP v18^96^ and of 18S rRNA ASVs (eukaryotes) using PR2 v5^97^. Consequently, the tables with taxonomic assignment, metadata, and ASV counts were uploaded into a Phyloseq object for filtering and statistical analyses^98^.

From the prokaryotic data (16S rRNA gene amplicons), reads that had only the Kingdom taxonomic level assigned were removed (29 ASVs), followed by reads of *Cutibacterium* (38 ASVs), Yersiniaceae (36 ASVs), Chloroplast (1186 ASVs), and tripletons (5,348 ASVs). From eukaryotic data (18S rRNA gene amplicon), reads that had only the Kingdom taxonomic level assigned were removed (881 ASVs), followed by reads of Metazoa (72 ASVs), Streptophyta (9 ASVs), and tripletons (8,393 ASVs). The ASV filtering approach followed the literature^64^. Similarly, we assigned taxonomy to Symbiodiniaceae reads (ITS2 rRNA) using the SymPortal pipeline^88^, and the sequences, taxonomy, and metadata tables were also uploaded into a Phyloseq object for filtering and statistical analyses with the removal of tripletons. Although there are only a few species of Symbiodiniaceae formally described, “Defining Intragenomic Variants’ (DIVs) are used instead of ASVs, while groups of DIVs that co-occur in the same coral colonies are categorized into ‘ITS2-type-profiles’^88^. When interpreting the DIVs, we took into account that the ITS2 rRNA gene has multiple copies^99^.

Statistical analyses were conducted in R Studio 1.3.1093^100^ using the packages vegan 2.6.4^101^ and ggplot2 v3.4.4^102^. Alpha diversity was estimated using the Shannon diversity metric. Statistical differences between the diversity of the two oceans were tested with ANOVA. Due to the non-parametric distribution of the data, differences in the diversity of each group of taxa between the ATO and IPO, and among global regions, were examined with the Kruskal-Wallis test followed by a post-hoc Dunn’s Test^103^. Samples were rarified and non-metric multidimensional scaling analyses (nMDS) of the Bray-Curtis dissimilarity distances were used to determine similarities in microbial communities due to sampling region and seawater temperature, and statistical differences were tested with pairwise Adonis 0.4.1^104^. The co-occurrence matrix of microbes associated with *Palythoa* samples was generated with SparCC^105^. Symbiont genera associated with >75% of *Palythoa* samples were considered core microbes^86^.

## Data and code availability

All raw sequencing reads are available under the NCBI BioProject accession number (XXXX). Tissue samples were deposited in the Fujukan Museum at the University of the Ryukyus, Okinawa, Japan (ESM Table 1). DNA vouchers of *Palythoa* samples used for sequencing are available in the laboratory of FH at the Okinawa Institute of Science and Technology. The full bioinformatics pipeline with all scripts is mirrored on the GitHub repositories https://github.com/SantosMEA and https://github.com/ECBSU.

## Acknowledgments

We thank the Sequencing (SQC) and Scientific Computing (SCDA) sections of the Okinawa Institute of Science and Technology for their great support. MEAS was supported by the JSPS postdoctoral fellowship 22KF0361 and the 2018 International Coral Reef Society Graduate Award. FH was supported by the JSPS Kakenhi grant 23K14256 and the HFSP Early Career Grant (RGEC29/2024). MEAS, JDR, MM, and FH were supported by the Aure grant 20D1000005. JDR was supported by JSPS Kakenhi grant 23H03821. HK was supported by JSPS Kakenhi grants 23KJ2206 and 19J12174. HBW was supported by the FRGS (FRGS/1/2021/WAB05/UKM/03/1), and GGPM (GGPM-2021-056). MVK thanks the support from the São Paulo Research Foundation (FAPESP #2021/06866-6) and the National Council for Scientific and Technological Development (CNPq #305274/2021-0). MM and AI were supported by Research Laboratory on Environmentally-conscious Developments and Technologies (E-code) at the National Institute of Advanced Industrial Science and Technology (AIST). RJT and AD were supported for work in Hawai’i by NSF-2048457 to RJT. Sampling in Brazil was supported by the grants SISBIOTA-Mar Network and CNPq 563276/2010-0 to MEAS and MVK.

