## Supplementary figures and images for "Coral microbiomes from the Atlantic and Indo-Pacific oceans have the same alpha diversity but different composition"

### Supplemental Figure 1

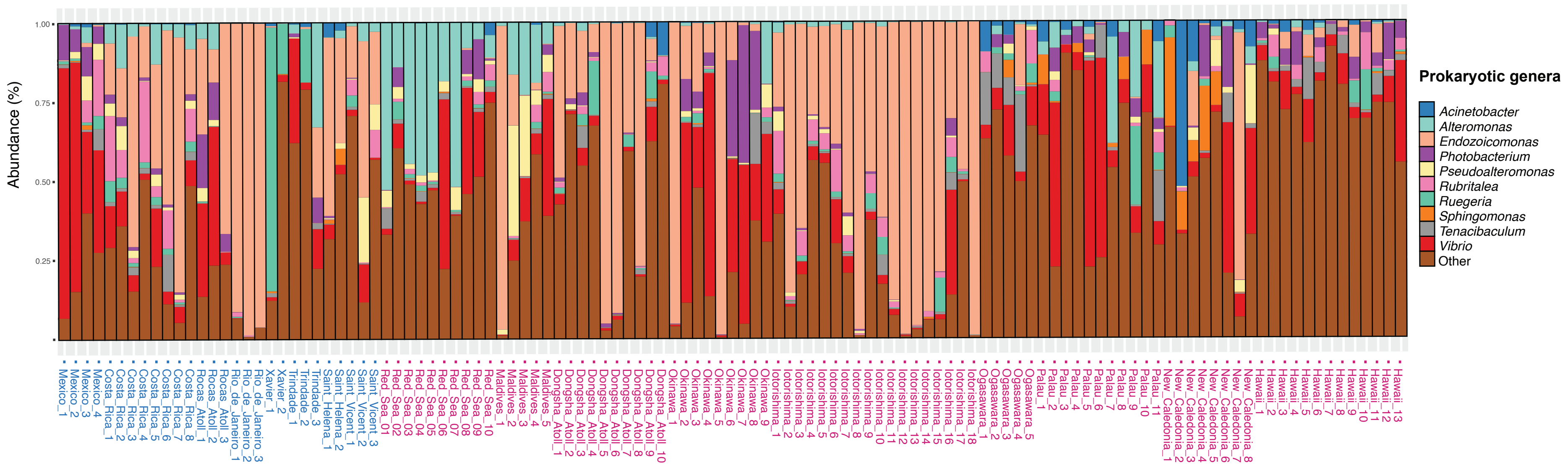
